# An optimized and validated workflow for developing stable producer cell lines with >99.99% assurance of clonality and high clone recovery

**DOI:** 10.1101/2022.12.16.520697

**Authors:** Julia Scherzinger, Daniel Türk, Fernando Aprile-Garcia

## Abstract

There is a constant pressure to reduce timelines in mammalian cell line development (CLD) for biotherapeutic protein production. Demonstration of clonal derivation of the generated cell lines is key for health authorities’ approval. To meet these regulatory and process-oriented demands, single-cell dispensers have become vital instruments for single-cell cloning. We conducted validation experiments with the UP.SIGHT (CYTENA GmbH) to determine this instrument’s single-cell dispensing efficiency (SCDE) and probability of clonal derivation (p(clonal)). Process optimization to maximize clone recovery with several cell lines was also performed, focusing on cloning media and plate type. With a SCDE >97%, p(clonal) >99.99% and clone recovery values of up to 80%, the data reported here support the notion that the UP.SIGHT covers all steps in the single-cell dispensing process with assurance of clonality and colony tracking, leading to faster and more efficient CLD workflows. This work also serves as a guideline for instrument validation and guidance towards process optimization.

## Introduction

Since the approval of monoclonal antibodies as biopharmaceuticals, a wide variety of biomolecules—from antibody-like molecules such as bispecific antibodies, Fc fusion proteins, and antibody fragments to viral vector particles for gene-therapy—and engineered cell lines for cell therapies have been developed.^1,2^ All these applications require stable genetic modification of cells. Cell line development (CLD) refers to the production of such a cell line, which enables production of the target biomolecule on a large scale or to obtain enough therapeutic cells. On the one hand, CLD is resource and time intensive since usually large numbers of clones must be screened for productivity, stability and product quality to find the best candidates for up-scaling and production. On the other hand, to ensure the quality, consistency and safety of these biologics, regulatory authorities require that new cell lines stem from a single-cell progenitor.^3^ Upon submission of a New drug application (NDA) or Investigational new drug (IND), organizations must provide evidence that their cell lines are clonally derived. The combination of these two factors creates a continuous need to improve CLD workflows for faster clone generation while providing assurance of clonal derivation.

The three most relevant performance metrics in single-cell cloning are single-cell dispensing efficiency (SCDE), probability of clonal derivation (p(clonal)) (also known as monoclonality assurance) and cloning efficiency. In other words, single-cell cloning methods of high cloning efficiency should be able to populate a microplate with single cells in most of the wells (as opposed to empty wells or wells containing more than one cell) in a process that assures a high probability that any given clone is truly derived from a single cell and that this process maintains cell viability well enough to provide a high percentage of wells with healthy growing colonies after single-cell dispensing.

Most of the methods used in mammalian CLD involve single-cell isolation with verification of the isolated cell in the well plate. Each method has its own advantages and disadvantages regarding ease of use, seeding and cloning efficiency as well as costs. The classical cloning method of limiting dilution is based on plating cells at a low density such that many wells contain only one cell, according to Poisson distribution.^4^ This method was adopted because of its ease of use and low cost. However, it has intrinsically low throughput and low efficiency since most of the wells remain empty or contain more than one cell. This leads to a time- and labor-consuming process because guaranteeing clonal derivation requires multiple rounds of serial subcloning.^5^ A widely used alternative method is fluorescence-activated cell sorting (FACS).^6^ The proportion of wells plated with only one cell can be much higher with FACS than with the limiting dilution method, but this method also has its own limitations. Sorting can be quite variable and obtaining high cloning efficiencies is usually challenging because of low clone recovery derived from the harsh isolation conditions the cells are exposed to in the sheath fluid during cell sorting including high voltage, high pressure and chemical stress. ^7^

A breakthrough method for single-cell deposition has been single-cell dispensing by droplet generation on demand. Compared to FACS and limiting dilution, this method achieves high cloning efficiencies by means of improved SCDE and assurance of clonal derivation.^8,9^ The single-cell dispensing technology employed in this method is based on proprietary, transparent microfluidic chips, real-time imaging technology and algorithms to sort and dispense single cells into 96-or 384-well microplates (CYTENA GmbH). To dispense single cells, the cell suspension is first loaded into a disposable cartridge that contains a microfluidic chip and then attached to the instrument’s dispenser. The dispenser’s piezo-electric plunger generates picoliter droplets of cell suspension on demand by displacing a small volume in the microfluidic chip without contacting the cell suspension. The exit of the cartridge nozzle is screened by an imaging system and the number, size and roundness of cells are tracked in real time. A pneumatic shutter situated below the nozzle is maintained in the open position, aspirating away droplets that are empty, contain multiple cells (negative events) or single cells that do not meet the chosen criteria until a positive single-cell event occurs. Droplets with single cells (positive events) are deposited into microplate wells pre-filled with media **(Figure 1A)**.^10,11^

**Figure 1.**
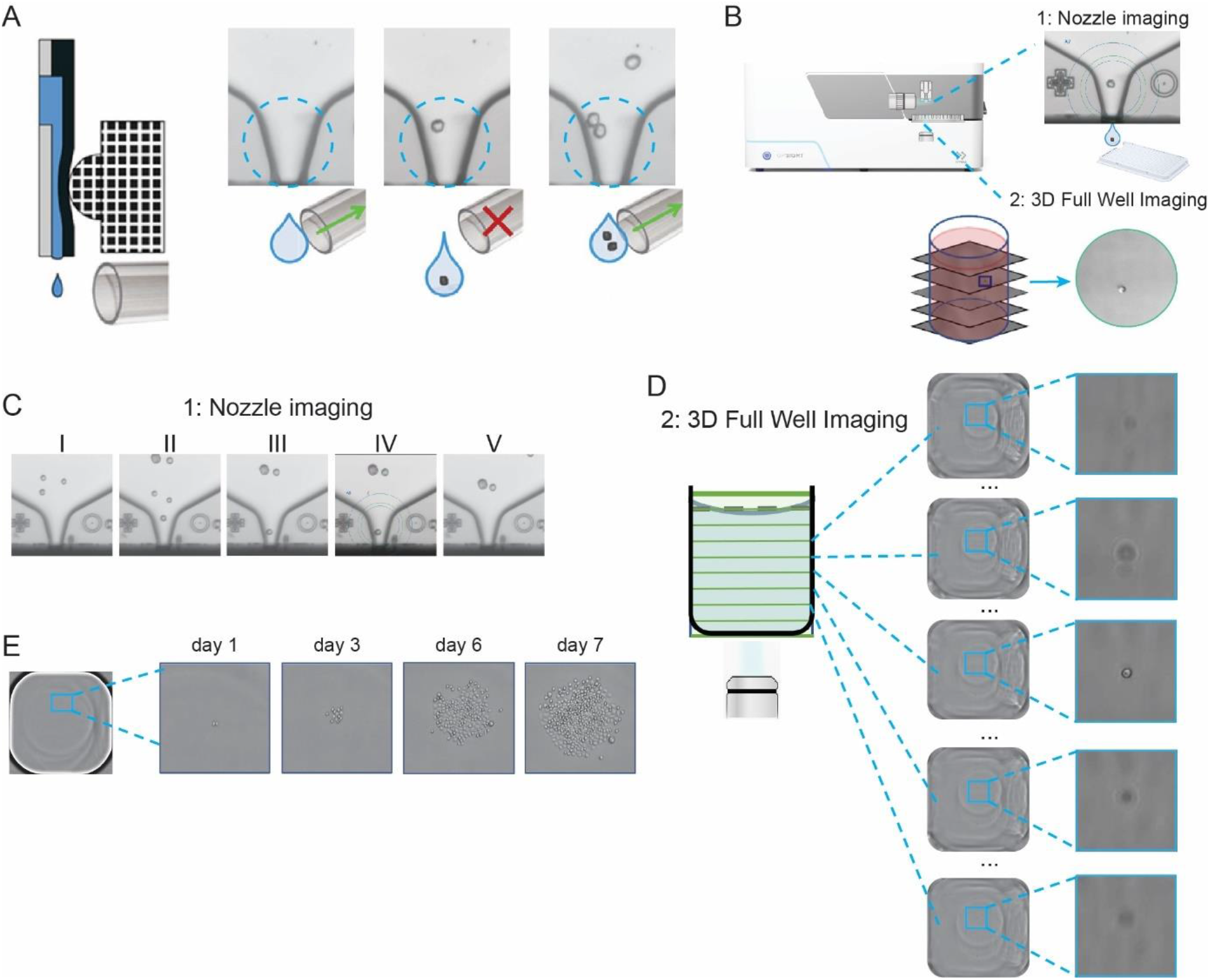
Working principle of the UP.SIGHT technology. A) Schematic of droplet generation on the dispenser chip (left), and shutter action to only allow droplets with single cells to pass unaffected and dispensed on the microplate (right). B) The two independent imaging systems of the UP.SIGHT: 1, a nozzle imaging system to assure that only droplets containing one cell are dispensed; and 2, a 3D Full Well Imaging system that generates a stack of images to confirm cell deposition. C) Example of a nozzle camera image series of single-cell deposition. A series of images is stored for each single-cell dispensing event: I, before the single-cell dispensing event; II, the cell enters region of interest (ROI); III, the cell is at the nozzle; IV, same image with image analysis overlay; and V, after the single-cell dispensing event to verify that the cell left the nozzle. D) Diagram of the 3D Full Well Imaging procedure with example images. A stack of images of an adjustable portion of the well height can be recorded directly after dispensing. E) The instrument can also be used as a classical plate imager to monitor colony growth. All the images generated can be imported into the C.STUDIO software for data analysis.

First generation single-cell dispensers do not provide visual confirmation of the single cell being deposited into the well of the destination plate. Therefore, plate imaging after dispensing is still required in limiting dilution or FACS methods. The combination of single-cell dispensing and plate imaging followed by image verification achieves very high levels of clonal derivation assurance (>99.99%).^12,13^ Nevertheless, plate imaging has its own limitations. It is usually difficult to detect single cells on the plate bottom. Sometimes the cell is not in the focal plane even after plate centrifugation or features of the plate itself such as edge effects, shadows or artifacts can hinder the ability to evaluate if a well contains a single cell. Those errors usually lead to “ghost wells,” where a colony grows without evidence of a single cell on Day 0 after dispensing and affects the overall assurance of clonal derivation.^14^ Because of this, the accuracy of the analysis is highly dependent on the plate’s quality and often cell staining is preferred to aid in cell detection.

Here, we employed a new instrument, the UP.SIGHT (CYTENA GmbH), to perform single-cell cloning. Compared to previous generations of single-cell dispensers, the UP.SIGHT provides two independent methods to document clonal derivation and plate imaging capabilities to enable colony tracking **(Figure 1B)**. For double assurance of clonal derivation, the device incorporates two independent imaging systems. The first is the classical nozzle camera that many cell dispensers have that images the dispensing nozzle to ensure only one cell is placed in each well **(Figure 1C)**. The second camera is located under the microplate and confirms that the single cells are inside the well with a new and innovative method called 3D Full Well Imaging. Directly after dispensing, the cell is detected and documented in the well volume by acquiring a z-stack of images, confirming that there is only one cell inside the well **(Figure 1D)**. This procedure eliminates concerns about “ghost wells” or artifacts and edge effects from well plates since it images the well volume and not the well bottom. Together, these two separate assurances of clonality provide verification of single-cell isolation during the dispensing process without requiring the extra procedures of centrifugation and bottom plate imaging. The images are saved and documented and can be used as supporting data for submissions to regulatory authorities. The optics module used for 3D Full Well Imaging can also image the plate bottom for colony tracking over time and clone characterization **(Figure 1E)**. As such, all the steps from single-cell dispensing with assurance of clonality and colony tracking are covered by the same instrument.

Here, we conducted experiments validating the UP.SIGHT’s capabilities. We provide experimental data showing high SCDE and (p(clonal)) by employing the UP.SIGHT in typical CLD campaigns. Furthermore, we provide data comparing the performance of culture media and well plates regarding clone recovery, raising the point of how important these two parameters are and call for their optimization for each cell line to be handled. These observations support the notion that the UP.SIGHT can replace—and for some parameters outperform—the current single-cell cloning gold standard of a single-cell dispenser coupled with a centrifuge and a plate imager.

## Results and discussion

### Single-cell dispensing efficiency

For the determination of SCDE and p(clonal), three independent experiments were performed with a total of 45 384-well plates dispensed (more than 17000 wells analyzed). Three UP.SIGHT instruments were employed in each experiment to rule out instrument-specific effects. Consistently, all three instruments performed in a similar way without noticeable deviations for all tested parameters (data not shown). CHOK1-GFP cells were dispensed into 384-well plates and after a brief centrifugation to bring the cells to the well bottom, the plates were imaged on a plate reader to detect how many cells were deposited in each well.

We first determined the SCDE of the UP.SIGHT. Out of more than 17000 wells analyzed, 97.82% contained one cell, 1.89% were empty and 0.29% of the wells had more than one cell—most often doublets **(Figure 2A)**. The UP.SIGHT’s nozzle images of the few empty wells and wells with more than one cell were visually examined to determine possible reasons for unsuccessful single-cell deposition. In most of the wells with more than one cell, it was easy to see the two cells on the nozzle images; the most frequent situation was one cell hiding behind another cell (28 events out of 48 doublets), followed by the second cell being very close to the nozzle wall and therefore not being detected as a cell (11 out of 48). There were only 5 cases where it was not possible to detect the two cells on the nozzle images **(Figure 2B)**. For empty wells, most of the time there was no apparent reason and the nozzle images showed one cell being dispensed. Only in some cases the nozzle images showed that the cell was not dispensed (38 out of 330) or that only debris was dispensed (4 out of 330) **(Figure 2B)**.

**Figure 2.**
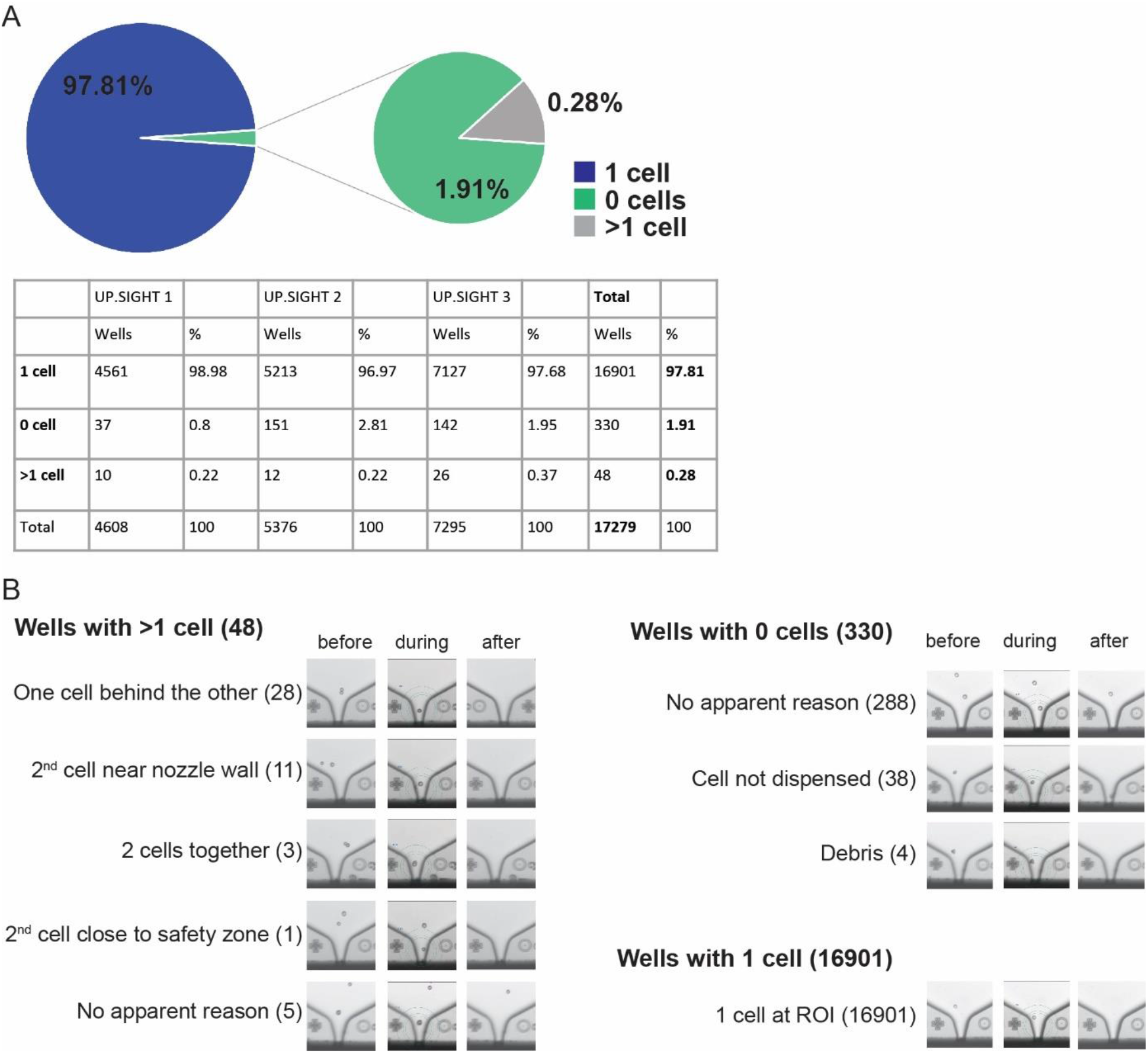
Determination of SCDE. A) SCDE was determined using three UP.SIGHT instruments each with three independent measurements, summing up 45 384-well plates dispensed. The pie chart and the table show the number of wells with either 0, 1 or 2 cells. B) Nozzle images from wells occupied by more than 1 cell (left), empty wells (top right) and 1 cell (bottom right) were grouped according to the image pattern observed. Three consecutive nozzle images are shown, one just before dispensing, one image during dispensing with the outline and safety area, and one image after dispensing.

All in all, these results show that the SCDE of the UP.SIGHT is high (>97.0 %) with a very low percentage of wells with more than one cell (<0.5 %). These are the wells that would be problematic during a typical CLD campaign because they would lead to cell lines of nonclonal origin. The process must assure with a high degree of confidence that there is a very low probability that these events remain undetected. For this reason, we then determined the probability of clonal derivation of the UP.SIGHT.

### Probability of clonal derivation

A production cell line derived from single-cell dispensing would be of nonclonal origin in case more than one cell was dispensed into the respective well. The probability of clonal derivation represents the likelihood that for any given well either the emerging cell line is of clonal origin, or if multiple cells got dispensed, they are successfully detected. On the UP.SIGHT, this probability is derived from the error rate of the nozzle images and of the well-stack images. These are considered to fail when they cannot recognize the unwanted multiple cells dispensed.

These error rates were determined by employing an experimental setup and statistical analysis previously reported for cloning by cell sorting coupled with plate imaging ^12,15^ (see Methods section for more details). The error rate of each imaging system was assessed by determining how many doublets (wells with more than one cell) were not detected by nozzle or 3D Full Well Imaging. Whenever a doublet was detected by the plate reader, the nozzle and the well-stack images were analyzed to evaluate if more than one cell could also be recognized on these images. Most of the time, it was easy to detect two cells on the images **(Figure 3A and 3B top panels for nozzle and well-stack images, respectively)**. If the two cells could not be discriminated on the nozzle or stack images, an error was called, accounting for the error rate of the respective imaging system **(Figure 3A and 3B bottom panels for nozzle and well-stack images, respectively)**.

**Figure 3.**
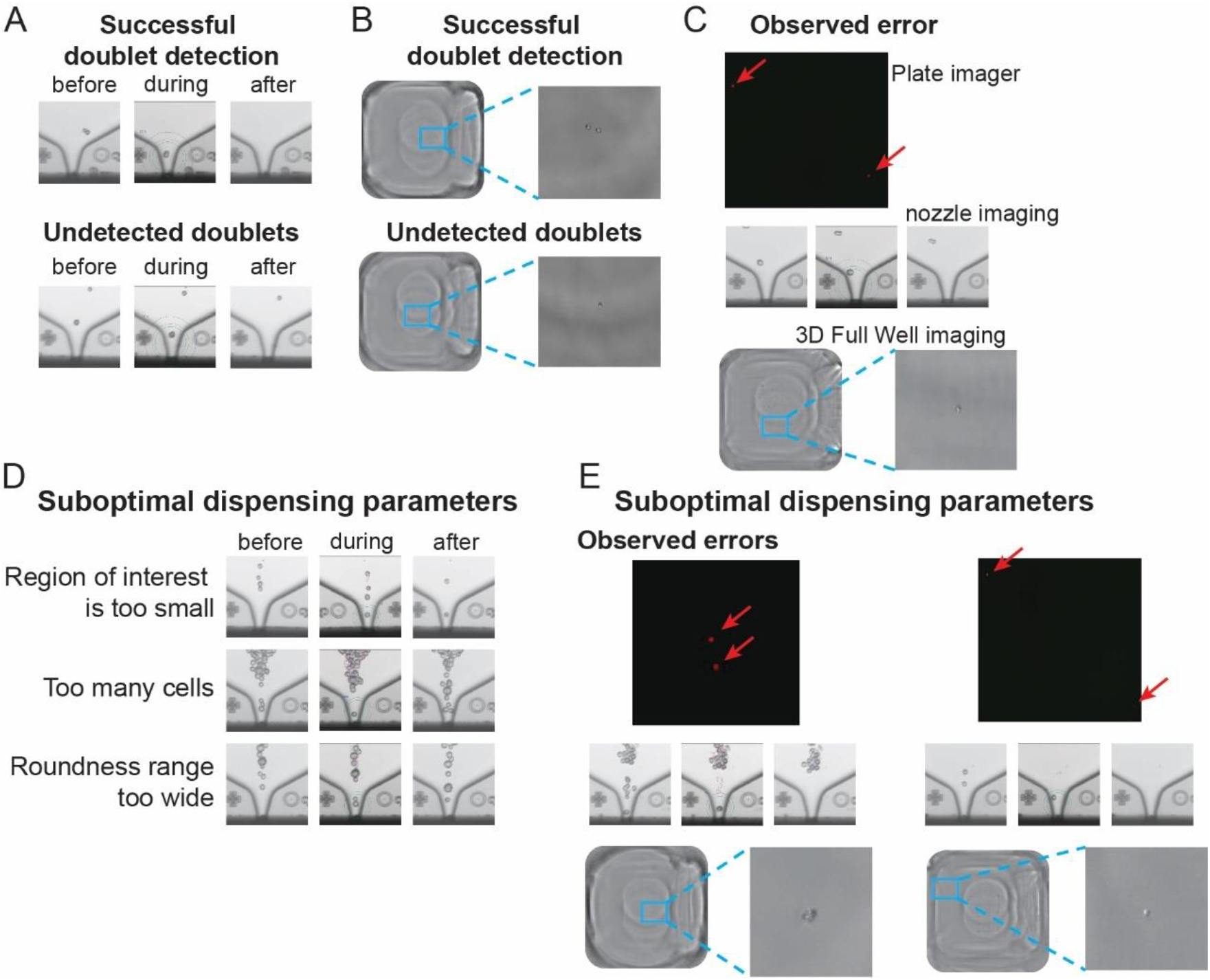
Probability of clonal derivation. A) Examples of nozzle images of wells with more than one cell dispensed like those shown in Figure 2B. Top example: it is easy to detect the doublet—the nozzle imaging system assists in determining this well is not clonally derived. Bottom example: the doublet is not readily detectable; the nozzle imaging system fails at detecting a doublet. This last example accounts for the nozzle detection error. B) Examples of individual stack images from the 3D Full Well Imaging system of wells with more than one cell dispensed. Top example: it is easy to detect the doublet, the 3D Full Well imaging system assists in determining this well is not clonally derived. Bottom example: the doublet is not readily detectable; the 3D Full Well Imaging system fails at detecting a doublet. This last example accounts for the 3D Full Well Imaging system detection error. C) Bottom plate, nozzle and well-stack images of the only well with more than one cell that remained undetected by both UP.SIGHT imaging systems after image verification. D) Nozzle images with suboptimal dispensing parameters that exemplify increased dispensing of droplets with more than one cell. E) Bottom plate, nozzle and well-stack images of the two wells with more than one cell that remained undetected by both UP.SIGHT imaging systems after image verification (dispensing with suboptimal parameters).

In the nine measurements performed (three instruments used over three independent experiments), five errors were found in the nozzle images among the 16,949 wells with at least one cell. With this observation, the statistical analysis based on the Wilson Method showed that at the upper limit of the 99% confidence interval, no more than 0.088% of the wells would contain more than one cell without detecting it upon image verification (see Methods section, Formula 1).

In the case of the 3D Full Well Imaging, 11 errors were found upon well-stack image examination. With this observation, the same statistical analysis showed that at the upper limit of the 99% confidence interval, no more than 0.139% of the wells would contain more than one cell.

Since the two imaging systems are independent, the probability of these two events occurring on the same well is the arithmetical product of both experimental errors:

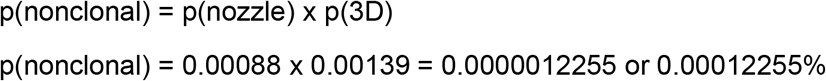

This leads to an overall probability of 0.00012255% that a cell line of non-clonal origin remains undetected after single-cell dispensing and image verification. This means that the probability of clonal derivation of the instrument, p(clonal), derived from the experimentally determined error rates is

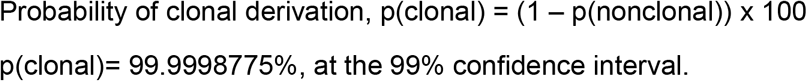

We then contrasted this *calculated error rate* (p(nonclonal)) with the *observed error rate* in this experiment. The observed error rate corresponded to the number of wells with doublets that remained undetected after nozzle and well-stack image examination. In this experiment, there was only one well for which this happened **(Figure 3C)**. Therefore, the observed error rate is 1/16949 = 0,000059, leading to a p(clonal)_obs_ of 99.9941%. This value should be taken with caution, however, since there was only one undetected error. More plates would need to be dispensed to find more errors for a more robust and accurate observed error rate.

Determining the instrument’s p(clonal) was intrinsically difficult because the doublet generation was very low (<0.5%, see **Figure 2A**), corresponding with a low error rate. Even with a very high number of plates dispensed (up to 45 in the results shown so far), only 48 doublets were generated and from those a very low number of errors were observed. This puts into question the statistical significance of the results obtained. To circumvent this, we looked for ways to artificially increase the number of doublets dispensed and again challenged the imaging systems. We purposely employed suboptimal dispensing parameters: higher cell suspension concentration, lower cell roundness threshold and smaller ROI. These settings increased the number of droplets that contained more than one cell and were not excluded by the nozzle imaging algorithm **(Figure 3D)**. Sixteen 384-well plates were dispensed, with 4,987 wells with at least one cell. Among those, 828 wells had more than one cell (16.6% doublet dispensing). From these, 24 errors were found by the nozzle and 13 errors were found by 3D Full Well Imaging upon image examination. As before, using Formula 1 at the upper limit of the 99% confidence interval, the nozzle detection error rate was not higher than 0.0081 and the 3D Full Well Imaging error rate was not higher than 0.0055, leading to a “suboptimal calculated error rate”:

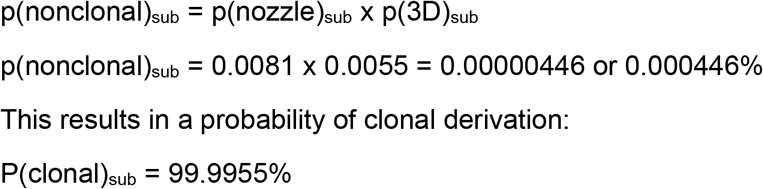

Even with suboptimal dispensing parameters that artificially increased the number of dispensed droplets containing more than one cell, the probability of clonal derivation was higher than 99.99%.

In this dataset, we detected instances in which multiple cells remained undetected. This would mean an observed error rate of 0.04% (2 errors in 4,987 wells). This observed error rate is an overestimation because of the increased generation of doublets caused by dispensing with suboptimal parameters. However, the overestimation is quantifiable: the first dataset (optimal dispensing parameters) had 0.28% of wells with more than one cell, whereas the second dataset (suboptimal dispensing parameters) had 16.6%. Therefore, the overestimation would be that of a 59.3 factor. If we apply this overestimation factor to the observed error rate with suboptimal dispensing parameters, it results in a 0.000000751 adjusted error, leading to a p(clonal)_sub-adj_ of 99.9993%.

To recapitulate, Table 1 shows the values obtained with the two datasets. Interestingly, p(clonal) and p(clonal)_sub-adj_ are very similar, suggesting that the actual p(clonal) of the UP.SIGHT should be very close to these values.

**Table 1.**
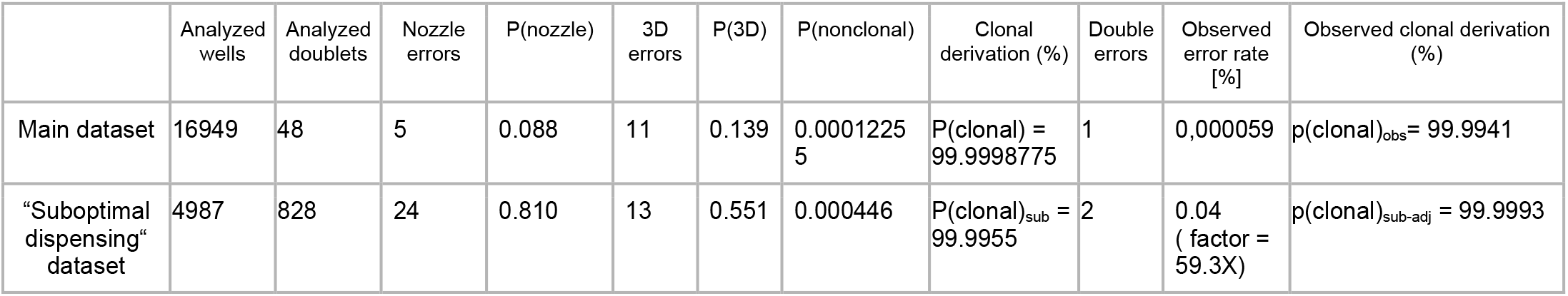
Results derived from the different datasets to estimate probability of clonal derivation using the UP.SIGHT.

In conclusion, the probability of clonal derivation is very high (>99.99%) and assures a single-cell cloning process where virtually all wells can be considered clonally derived after visual examination of nozzle and well-stack images.

### Clone recovery optimization: influence of culture media and plates

The last important metric for single-cell cloning is clone recovery. The culture conditions of the single cells after dispensing are key to a high recovery rate. Cloning media and the cell culture plates used can influence the growth of different cell lines. To further study these variables, we conducted several single-cell dispensing experiments where plates were imaged with the UP.SIGHT 14 days later to track colony growth and clone recovery.

We used a CHO-derived cell line overexpressing the human IgG1 monoclonal antibody Trastuzumab, CHOK1-Her2, since it is a good example of a typical CLD application: human monoclonal antibody production. We first cultivated cells in two cultivation media designed to support growth of CHOK1 cells. Each culture was then dispensed into three different cloning media **(Figure 4A)**. As a general trend, cells cultivated in CHOlean (green bars) led to slightly better clone recovery values than those cells cultivated in CDCHO (blue bars). Cultivation in CHOlean also led to a more homogeneous recovery regardless of the cloning medium employed. Regarding cloning media, CDCHO led to better clone recovery than the other two media tested **(Figure 4A)**. These data suggest that a careful selection of cultivation and cloning media can improve clone recovery. We further focused on cloning medium optimization. We compared three commercially available cloning media designed to support the growth of CHO single cells. The performance was variable, from little more than 10% recovery to up to 60% **(Figure 4B, blue bars)**. In addition, it is known that cloning medium supplementation can boost single cell survival.^16,17^ One typical supplementation strategy is the use of conditioned medium; i.e., supplement fresh medium with medium used to grow cells and thus containing growth factors, signaling molecules, etc.^18^ When the cloning medium was supplemented with conditioned medium at an 80:20 ratio, the clone recovery in each case improved substantially **(Figure 4B, green bars)**. In addition, the large differences observed between the raw media tested was buffered by the supplementation with conditioned medium, obtaining more homogeneous clone recovery values from 60% to 80%.

**Figure 4.**
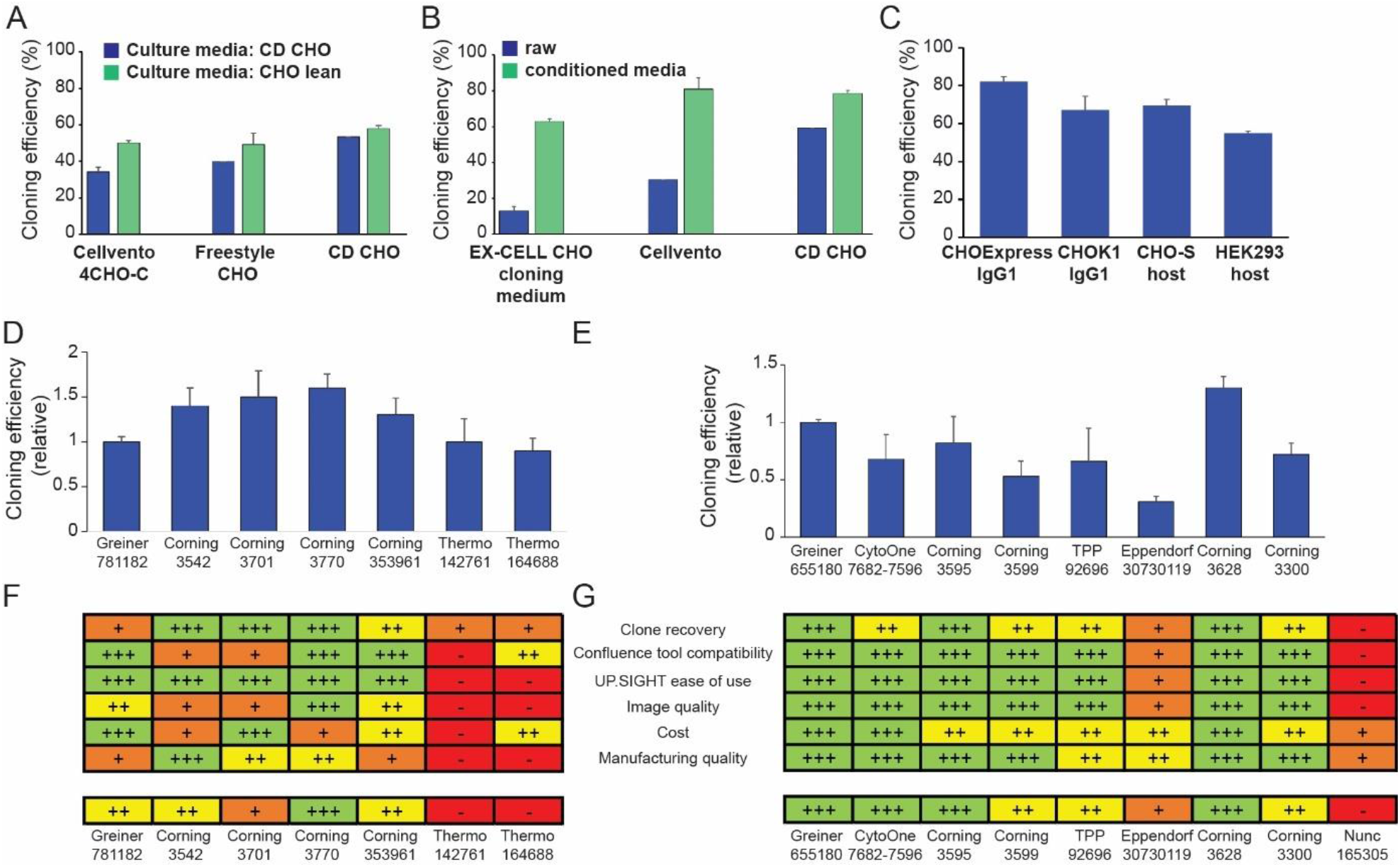
Clone recovery optimization. A) Cultivation and cloning medium optimization for single-cell dispensing of CHOK1 cells. Cells were grown in two different culture media and dispensed in three cloning media each. Clone recovery values are expressed as percentage of wells occupied by colonies on 384-well plates after 14 days of cloning. N=2, error bars: s.e.m. B) Cloning medium optimization for single-cell dispensing of CHOK1 cells and effect of conditioned medium. Cells were dispensed in three cloning media, supplemented or not with conditioned medium. Clone recovery values are expressed as percentage of wells occupied by colonies on 384-well plates after 14 days of cloning. N=2, error bars: s.e.m. C) Example of typical clone recovery values which can be obtained after medium optimization for several examples of suspension cells. N=3 to 6, error bars: s.e.m. D) and E) Clone recovery values after dispensing CHOK1 cells in different 384-or 96-well plates, respectively. Values are relative to the reference plate from Greiner, first column on the left. F) and G) Evaluation of several parameters relevant to plate usability on the UP.SIGHT for 384- and 96-well plates, respectively. Results are shown using a qualitative, color-coded scheme. The tables provide a quick reference for choosing the right plate according to each experimental requirement, including a summary for overall usability assessment.

We then optimized cell culture conditions for several suspension cell lines, typically employed in CLD campaigns for therapeutic protein production or gene therapy applications. This included CHO-derived cell lines – some producing human IgG1s – and a HEK293 suspension host cell line. We established the best cloning medium for each cell line in terms of clone recovery (data not shown). After optimization, consistent high clone recovery values were obtained after single-cell dispensing **(Figure 4C)**. All in all, the data show that a careful selection of cloning medium and supplements as well as cultivation medium is important to optimize clone recovery after single-cell dispensing. The optimal conditions will be cell line-specific.

We also examined the influence of cell culture plates on clone recovery. The evaluation of a selection of plates led to the conclusion that clone recovery is affected by the surface employed during single-cell cloning. Recovery values were highly variable with 384- and 96-well plates from different manufacturers **(Figure 4D and E, respectively)**. Other factors should also be considered when selecting plate types for single-cell cloning. The design of the plates and their lids are known to influence evaporation rates of media, particularly in the outer, more exposed wells. This factor will impact clone recovery and cell health. Another important aspect is plate imaging; plate and well geometry can have an impact on imager compatibility and image quality. Also, manufacturing quality is important to minimize wells with scratches or artifacts on their surface and to have well rows and columns properly aligned to support automated plate imaging. Finally, cost evaluation should be considered. These features were evaluated, and qualitative, color-coded tables were composed **(Figure 4F for 384-well plates and Figure 4G for 96-well plates)**. This becomes a helpful resource when selecting the right type of plate for single-cell cloning experiments using the UP.SIGHT.

## Conclusions

In this work, we measured the performance of the UP.SIGHT single-cell dispenser for CLD applications. Single-cell dispensing efficiency, clonal derivation assurance and cloning efficiency are key metrics for robust CLD processes. Here, we have demonstrated that single-cell dispensing with the UP.SIGHT confers extremely high SCDE and assurance of clonal derivation. The results show that for every 384-well plate dispensed, only a few wells will remain empty and not more than two wells should be occupied by more than one cell. To assure clonal derivation, the combination of the nozzle images and well-stack images allowed the detection of doublets in almost every situation after image evaluation. We have also provided experimental data covering optimization of clone recovery. We have shown how important a careful culture medium and plate selection can be to maximize colony outgrowth after single-cell dispensing, highlighting the point that this is usually cell line-dependent.

Given the fact that the UP.SIGHT can also image the plate bottom for colony tracking and clone characterization over time, all the steps from single-cell dispensing with assurance of clonality and colony tracking can be covered by the same instrument. This should not only result in a faster and more efficient cloning workflow, but also in better documentation for improved quality of the final biological product.

The methods and results shown in this work should serve as a guideline for novel UP.SIGHT users that might require instrument validation and guidance towards process optimization.

## Materials and Methods

### Cell culture

All cell lines used in this work were cultivated in cell culture incubators at 37°C and 5% CO_2_. GFP-overexpressing CHO-K1 cells (Linterna CHO-K1, Innoprot) were cultivated in DMEM (ThermoFisher Scientific) supplemented with 1% Penicillin/Streptomycin, 1% Glutamax, 1% NEAA, 10% FBS. The following cell lines were cultivated in 125 mL shaken flasks at 130 rpm on a 25 mm diameter orbital shaker: IgG1 Trastuzumab overexpressing CHOK1 cells (CHOK1-Her2, a kind gift from Professor Dr. Nicole Borth from the Institute of animal cell technology and systems biology BOKU, Vienna, Austria) were grown either in CD CHO medium (ThermoFisher Scientific) or CHOlean (Xell AG) with 1% Penicillin/Streptomycin and 1% Glutamax; CHO-S cells (ThermoFisher Scientific) were grown in Freestyle CHO expression medium (ThermoFisher Scientific); suspension HEK293 host cell line (Freestyle 293-F, ThermoFisher Scientific) was grown in Freestyle 293 expression medium (ThermoFisher Scientific). CHOExpress® cells were cultivated according to supplier’s recommendation (Excellgene SA).

### Single-cell dispensing

Cells were cultivated for at least three passages after thawing before single-cell dispensing. 24 hours prior to dispensing, the cell culture was adjusted to 1×10^6^ cells/mL in fresh medium to assure high viability and an exponential growth phase of the culture at the time of dispensing. Only cultures with a viability higher than 90% were used for single-cell dispensing. Cells were harvested, washed once with PBS (room temperature centrifugation, 3min at 300g), counted and resuspended in PBS to an end concentration of 1×10^6^ cells/mL. The EASY.ON Cartridge (CYTENA GmbH) was filled with 100 μL of cell suspension and loaded on the UP.SIGHT (CYTENA GmbH) according to the manufacturer’s instructions (“2022 UP.SIGHT practical guide_R3”, CYTENA). Cells were dispensed in 96-or 384-well plates and either imaged on a plate imager (SCDE and p(clonal) experiments) or incubated for 14 days to assess colony growth (clone recovery optimization experiments).

To determine SCDE and p(clonal) of the UP.SIGHT, GFP-CHO cells were dispensed in 384-well plates filled with 50 μL of DMEM. The UP.SIGHT was setup with cartridge agitation on 20 μL volume at 10 repetitions per minute, and laser turned on at 21 mW of power and 1 ms of exposure. Cells were sorted according to roundness (0.6 to 1), size (from 15 to 25 μm diameter) and fluorescence (from 90 to 255 intensity values). A half-stack of images was obtained with the 3D Full Well Imaging system. Manufacturer’s instructions (“2022 UP.SIGHT practical guide_R3”, CYTENA) were followed to configure the imager settings to obtain half-stacks of 2.84 mm in eight (50 images). These settings assured that the dispensed cells would be captured by at least one of the images of the configured half-stack.

### Suboptimal dispensing conditions

To artificially generate an increased number of dispensed doublets for the determination of p(clonal)_sub,_ we employed suboptimal dispensing parameters. These settings would never be used in a real setting but were useful for the purposes of forcing the instrument to dispense more than one cell. We employed a more concentrated cell suspension at 2×10^6^ cells/ml and switched the cell focusing function on in order to have a cartridge saturated with cells. The cell parameters were loosened: roundness from 0.2 to 1 and size of 0.15 μm diameter and higher. The safety zone around the ROI was eliminated and the inner ROI was reduced to a size smaller than an actual droplet. All these changes led to increased doublet dispensing **(Figure 3D)**.

### Clone recovery experiments

Cells were dispensed into the indicated plate and media types. After dispensing, plates were transferred to the incubator, where they were kept for 14 days to allow single-cell recovery and colony growth. On day 7, fresh cloning medium was added (50 μL or 100 μL for 384- and 96-well plates), resulting in 100 or 200 μL total volume, respectively. At day 14, well bottom images were acquired with the UP.SIGHT. Images were taken with the dispenser flapped out for better image quality. For every plate type used, the correct focus was manually adjusted and the “autofocus during run” option enabled. The best well plate adapter frame was selected and general compatibility with the imager was evaluated (plate geometry calibration, plate fitting the imaging stage). Images were also analyzed with respect to manufacturing defects such as bottom scratches, stains and features hindering clear images.

The acquired images were automatically analyzed by the confluence tool in the C.STUDIO software (version 1.1.3). This tool employs a colony-detection algorithm to quantify the surface of a well occupied by the colony. A 0.5 mm^2^ surface threshold was employed in these experiments to calculate cloning efficiency values. Results are expressed as % of total wells occupied with colonies larger than the threshold size.

### Bottom imaging with a conventional plate imager

After dispensing, the plates were spun down at 50 g for two minutes and then imaged on a plate imager (NyOne, SYNENTEC). The FASCC plugin was used to determine the number of GFP-fluorescent cells in each well. A careful setup and validation of this tool (i.e., adjusted imaging and detection parameters to recognize all the cells without background or false positives) was required (data not shown). After that, almost 100% of the wells that reportedly contained one single cell were indeed correct. In contrast, the images corresponding to wells reportedly containing a cell number other than one were inspected manually to discard NyOne software detection errors. These corrected values were used to calculate SCDE and p(clonal).

### Determination of probability of clonal derivation

After manual correction of the NyOne evaluation, a list of wells containing doublets was assembled for each dataset. The nozzle images and the well-stack images generated by the UP.SIGHT for these specific wells were evaluated to find doublets that were not readily apparent, therefore accounting for the error rate. To account for experimental variability and sample size (i.e., the number of evaluated wells), the probability of error of each imaging system—(p(nozzle) and p(3D), respectively—was calculated with a one-sided upper 99% confidence interval using the Wilson method:^19^

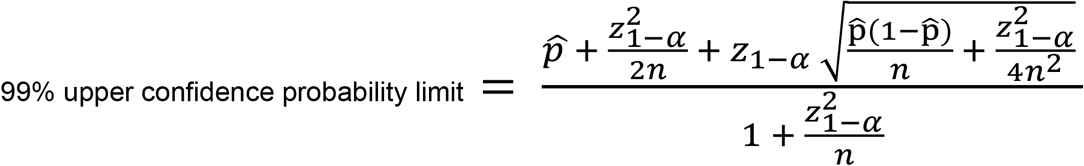

**Formula 1**. Where 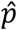 is the observed proportion of wells with more than one cell that are not readily detected on the images. *z*_1−*α*_ is the 1-ath percentile of the standard normal distribution (*z*_1−*α*_ =2.58) and *n* is the total number of wells with at least one cell determined by plate imaging.

The probability of error of the instrument p(nonclonal) is the arithmetical product of both probability values, p(nonclonal) = p(nozzle) x p(3D).

The probability of clonal derivation, p(clonal) expressed as a percentage, results from: p(clonal) = (1 – p(nonclonal)) x 100

## Acknowledgments

The authors would like to thank Katrin Raute and Julian Riba (CYTENA), and Stéphanie Anchisi and Benjamin Moritz (Excellgene SA) for critical reading of the manuscript.

## References

1. Chen YH, Pallant C, Sampson CJ, et al. Rapid lentiviral vector producer cell line generation using a single DNA construct. Molecular Therapy--Methods & Clinical Development. 2020; 19: 47–57.

2. Flahou C, Morishima T, Takizawa H, Sugimoto N. Fit-for-all iPSC-derived cell therapies and their evaluation in humanized mice with NK cell immunity. Frontiers in Immunology. 2021; 12.

3. Plavsic, M. Q5D Derivation and Characterization of Cell Substrates Used for Production of Biotechnological/Biological Products. ICH Quality Guidelines. 2017: 375–393.

4. Coller HA, Coller BS. [37] Poisson statistical analysis of repetitive subcloning by the limiting dilution technique as a way of assessing hybridoma monoclonality. Methods in Enzymology. 1986; 121: 412–417.

5. Underwood PA, Bean PA. Hazards of the limiting-dilution method of cloning hybridomas. Journal of Immunological Methods. 1988; 107(1): 119–128.

6. Herzenberg LA, Parks D, Sahaf B, et al. The history and future of the fluorescence activated cell sorter and flow cytometry: a view from Stanford. Clinical Chemistry. 2002; 48: 1819–1827.

7. Yoshimoto N, Kida A, Jie X, et al. (2013). An automated system for high-throughput single cell-based breeding. Science Reports. 2013 31 3, 1–9.

8. Bhagat AAS, Bow H, Hou HW, et al. Microfluidics for cell separation. Medical & Biological Engineering & Computing. 2010; 48: 999–1014.

9. Yusof A, Keegan H, Spillane CD, et al. Inkjet-like printing of single-cells. Lab on a Chip. 2011; 11(14): 2447–2454.

10. Gross A, Schoendube J, Zimmermann S, et al. Technologies for single-cell isolation. International Journal of Molecular Science. 2015; 16(8): 16897–16919.

11. Gross A, Schöndube J, Niekrawitz S, et al. Single-cell printer: automated, on demand, and label free. Journal of Laboratory Automation. 2013; 18(6): 504–518.

12. Evans K, Albanetti T, Venkat R, et al. Assurance of monoclonality in one round of cloning through cell sorting for single cell deposition coupled with high resolution cell imaging. Biotechnology Progress. 2015; 31(5): 1172–1178.

13. Yim M, Shaw D. Achieving greater efficiency and higher confidence in single-cell cloning by combining cell printing and plate imaging technologies. Biotechnology Progress. 2018; 34(6): 1454–1459.

14. Chen C, Le K, Le H, et al. (2020). Methods for estimating the probability of clonality in cell line development. Biotechnology Journal. 2020; 15(2).

15. Pybus LP, Kalsi D, Matthews JT, et al. Coupling picodroplet microfluidics with plate imaging for the rapid creation of biomanufacturing suitable cell lines with high probability and improved multi-step assurance of monoclonality. Biotechnology Journal. 2022; 17(1).

16. Ritacco FV, Wu Y, Khetan A. Cell culture media for recombinant protein expression in Chinese hamster ovary (CHO) cells: history, key components, and optimization strategies. Biotechnology Progress. 2018; 34(6): 1407–1426.

17. Pérez-Rodriguez S, Ramírez-Lira MdJ, Trujillo-Roldán MA, Valdez-Cruz NA. Nutrient supplementation strategy improves cell concentration and longevity, monoclonal antibody production and lactate metabolism of Chinese hamster ovary cells. Bioengineered. 2020; 11(1): 463–471.

18. Lim UM, Yap MGS, Lim YP, Goh LT, Ng SK. Identification of autocrine growth factors secreted by CHO cells for applications in single-cell cloning media. Journal of Proteome Research. 2013; 12(7): 3496–3510.

19. Agresti A. Categorical Data Analysis. 2nd ed. John Wiley & Sons, Inc; 2002

